# An RNAi screen in human cell lines reveals conserved DNA damage repair pathways that mitigate formaldehyde sensitivity

**DOI:** 10.1101/310730

**Authors:** Eleonora Juarez, Nyasha Chambwe, Weiliang Tang, Asia D. Mitchell, Nichole Owen, Anuradha Kumari, Raymond J. Monnat, Amanda K. McCullough

## Abstract

Formaldehyde is a ubiquitous DNA damaging agent, with human exposures occuring from both exogenous and endogenous sources. Formaldehyde can also form DNA-protein crosslinks and is representative of other such DNA damaging agents including ionizing radiation, metals, aldehydes, chemotherapeutics, and cigarette smoke. In order to identify genetic determinants of cell proliferation in response to continuous formaldehyde exposure, we quantified cell proliferation after siRNA-depletion of a comprehensive array of over 300 genes representing all of the major DNA damage response pathways. Three unrelated human cell lines (SW480, U-2 OS and GM00639) were used to identify common or cell line-specific mechanisms. Four cellular pathways were determined to mitigate formaldehyde toxicity in all three cell lines: homologous recombination, double-strand break repair, ionizing radiation response, and DNA replication. Differences between cell lines were further investigated by using exome sequencing and Cancer Cell Line Encyclopedia genomic data. Our results reveal major genetic determinants of formaldehyde toxicity in human cells and provide evidence for the conservation of these formaldehyde responses between human and budding yeast.

## INTRODUCTION

Endogenous and environmental exposures to formaldehyde are well-correlated with an increased risk of cancer, asthma, and other diseases (1–4). Formaldehyde is produced endogenously as an essential component of human cellular metabolism including one carbon pool, amino acid and alcohol metabolism, lipid peroxidation, and P450-dependent demethylation (1). In humans, the steady state level of formaldehyde in whole blood or plasma is remarkably high, ranging between 22–87 μM (5–7). However, these reported blood levels may be an overestimate in light of a recent study that failed to detect any endogenous formaldehyde-serum albumin adducts (summarized in (8)). Some cell lineages, e.g., hematopoietic stem cells, may receive higher exposure via lineage-specific generation of formaldehyde from histone demethylation as part of chromatin remodeling during differentiation (9,10). Human tumor cells can be stimulated to generate formaldehyde in response to treatment with widely used chemotherapeutic agents, such as anthracyclines (11,12). Environmental exposure to formaldehyde includes occupational exposures and sources such as automobile exhaust, cigarettes and e-cigarettes, cosmetic products, forest fires, and manufactured wood products (13–19).

The genotoxicity and ubiquitous nature of formaldehyde exposure have driven efforts to better understand the cellular pathways that mitigate formaldehyde toxicity. Specific cellular processes reported to promote cell survival include Nucleotide Excision Repair (NER) (20–22), proteasomal degradation (23), metalloproteases (24,25), the Fanconi Anemia pathway (26–29), and Homologous Recombination (HR) (20,22,30,31). We and others have also shown that formaldehyde can perturb the cell cycle and alter gene expression (21,30,32–35). In order to systematically analyze the role of DNA damage repair (DDR) pathways in modulating formaldehyde toxicity, we selectively depleted each of 320 genes representing key members of the major DNA damage repair, cell cycle, and mitotic cell division pathways. Gene depletion was followed by quantification of cell proliferation suppression as a function of formaldehyde dose. The resulting library is referred to herein as the 320 DDR (320 gene DNA damage response) library. This library was used to screen three well-characterized, though otherwise unrelated, human cell lines to identify genes that modified cell proliferation in response to chronic formaldehyde exposure. The cell lines, GM00639, SW480, and U-2 OS, were derived respectively from primary human fibroblasts, an epithelial adenocarcinoma, and an osteosarcoma, and were chosen due to their wide-spread use in genotoxic studies (30,36–38).

Our results identify four pathways that strongly influenced cellular proliferation after formaldehyde exposure: HR, double-strand break (DSB) repair, ionizing radiation (IR) response, and DNA replication. These results are concordant with prior work in budding yeast where we, and others, have identified genes and pathways important for suppression of cell proliferation following formaldehyde exposure (20,22). Thus, our work broadens the foundation from which to understand the mechanistic determinants of formaldehyde toxicity in human cells.

## MATERIALS AND METHODS

### Cell lines

Three well-characterized human cell lines were used for our screens: GM00639, SW480, and U-2 OS. GM00639 is a widely used human fibroblast cell line that was derived by SV40 transformation of primary fibroblasts from an 8-year-old galactosemic male (39). SW480 is an epithelial colorectal adenocarcinoma cell line (40). U-2 OS is an osteosarcoma cell line derived from a 15-year-old female (41). GM00639 and U-2 OS cells were kind gifts from Dr. Robb Moses and SW480 from Dr. Owen McCarty (both at Oregon Health & Science University). Cells were grown in DMEM supplemented with 10% fetal bovine serum and antibiotic/antimycotic (penicillin, streptomycin, and Amphotericin B, Gibco) at 37°C in a humidified, 5% CO2 ambient oxygen incubator.

### Genomic analyses

Coding region variant calls for the SW480 and U-2 OS cell lines were downloaded from the Cancer Cell Line Encyclopedia (CCLE) Project web-based data portal (42) (merged variant maf file: CCLE_DepMap_18Q1_maf_20180207.txt). Briefly, this merged file integrates variant calls from CCLE whole genome and exome sequencing (WGS, WES), CCLE RNA sequencing (43), and WES generated by the Sanger Institute as part of the COSMIC project and is available at the European Genome-phenome Archive (EGAD00001001039). We filtered out variant calls sourced from the RNA-Seq data alone, and those with low alternate allele counts from any of the WES/WGS platforms.

To generate comparable data for GM00639, we carried out exome capture and sequencing of a clonal derivative, GM639-CC1 (39,44). Exome capture was performed using Nimblegen SeqCap EZ HGSC VCRome kit (V2), with sequencing performed on an Illumina HiSeq to generate 200 bp read lengths. Sequence reads were aligned to the hg19 human reference genome using BWA (BWA-MEM) (45). We applied GATK (46) indel realignment and base quality score recalibration according to GATK best practices recommendations (47,48). Variant calling by GATK UnifiedGenotyper was restricted to ±1000 bp around the capture regions (VariantFiltration module). Median read depth was 109 and prior to filtering variants were annotated using in-house scripts including the ANNOVAR pipeline (49). We filtered variants using six criteria: coverage ≥ 30, GATK hard filter pass, read quality ≥ 20 (pred-based), read depth ≥15, variant allele frequency ≥15%, and exclusion of common variants using a population minor allele frequency threshold of 1%. Population allele frequencies were mined from the Exome Variant Server, NHLBI GO Exome Sequencing Project (ESP) (http://evs.gs.washington.edu/EVS/) [ESP6500], 1000 Genomes Project (phase three) (50), The Exome Aggregation Consortium (ExAC) (51) and two internal exome collections with 650 and 945 exomes, respectively.

To enable joint analysis across three cell lines, we merged and re-annotated variant calls using the Oncotator (52) pipeline, generated a unified variant annotation and then performed additional filtering to exclude synonymous mutations.

Copy number data for the SW480 and U-2 OS cell lines were downloaded from the CCLE web portal (42). Gene-level copy number estimates (normalized log ratios) were inferred from genome-wide Affymetrix SNP6.0 array data as previously described (42). Using a strategy similar to that used by Kim *et al*. (53), we determined genes that were amplified or deleted using a threshold of ± 0.7 to identify approximate two-fold changes for mean segment values that estimate copy number.

### Dose-dependent formaldehyde-induced suppression of cell proliferation

We determined dose-dependent formaldehyde toxicity by quantifying cell proliferation suppression after treating cells continuously with formaldehyde for 5 days. Assays were performed in triplicate in 96-well plates: cells were plated at sub-confluent density, allowed to attach overnight, and then treated with formaldehyde (Fisher Scientific) at indicated doses. Viable cell number was assessed on Day 5 using Cell Titer-Glo^®^ (CTG) following the manufacturer’s instructions. CTG assesses viable cell numbers by quantifying ATP generated by metabolically active cells. Briefly, 100 μl of CTG reagent was added to each well prior to mixing the plate on a shaker for 10–15 min. Luminescence output for each well was quantified on a Tecan plate reader (Infinite M200). The suppression of cell proliferation as a function of formaldehyde dose (expressed as GI_25–75_) was calculated for each cell line using Graph Pad Prism 7 software (La Jolla, CA, USA) with a sigmoidal, 4PL log curve fit.

### RNAi screens

A custom-designed RNAi library was used that consisted of a pool of four siRNAs targeting each of 320 genes representing all major DNA repair pathways with additional genes involved in the DNA damage response, cell cycle regulation, and mitotic cell division (54). All siRNAs were synthesized on a 0.25 μmol scale, and then each gene-specific set of 4 siRNAs was pooled in a single master plate well (Qiagen). Each gene-specific pool was tested in triplicate on separate plates to establish experimental variability, statistical validity, and to identify potential batch effects. The full 320 DDR RNAi library and sequences of gene-specific and control siRNAs are provided in Supplementary Table 1 of Kehrli *et al* 2016 (54). This siRNA library is available for both academic and commercial use as the 320 DDR (DNA Damage Repair) Library through the University of Washington Quellos High Throughput Screening Core (http://depts.washington.edu/iscrm/quellos/rnai-screens).

RNAi screens for gene depletion and formaldehyde dose-dependent suppression of cell proliferation were performed in 384-well format on the Quellos High Throughput Screening Core platform. Transfection conditions for siRNA were first optimized for each cell line using an siRNA that targets the *KIF11* kinesin family member 11 gene, whose encoded protein arrests cells in mitosis. We used as a minimum threshold, the loss of at least 50% cell viability with <25% absolute deviation upon *KIF11* siRNA transfection versus a control siRNA or mock-transfected cells (cells plus media and Optimem only). Transfection optimization and screens were performed using Lipofectamine RNAiMAX (Invitrogen) according to the manufacturer’s instructions. Mock and non-targeting universal siRNAs were used in addition to the *KIF11* siRNA as transfection and RNAi pathway-dependent controls.

The *KIF11* positive control was siRNA SASI_Hs01_00161689, and the siRNA negative control was the MISSION siRNA SIC001 Universal Control #1, both purchased from Sigma-Aldrich. All reagent conditions were statistically evaluated using a simple Z-factor score (all scores ≥ 0.5) to determine differences and variability among replicates, and to identify optimal transfection and treatment conditions for each cell line.

We expected a subset of siRNAs to exert cytotoxic effects and identified these by determining the effect of siRNA transfection alone on cell proliferation using ≥ 20% cell viability (≤ 80% cell death) as a cut-off to remain within assay detection limits and biological plausibility (data not shown). Concentrations of formaldehyde leading to no (*i.e*., GI0) or 10–90% (*i.e*., GI10-90) proliferation suppression were determined for each cell line by formaldehyde titration over a 0 – 100 μM range. We verified reproducibility across replicates for each formaldehyde dose, then calculated the standard deviation to determine the variation across sample replicates. Z-scores for the standard deviation were calculated using the equation,

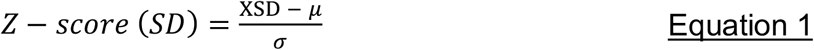

where *μ* is the mean of the standard deviations and σ the standard deviation of the mean standard deviations. This determined acceptable reproducibility across replicates for each formaldehyde dose in our assays. Genes with Z-scores ≥ +2.0, or those for which an untreated data point did not meet the quality minimums outlined above, were excluded.

These results were used to design RNAi screens in which cells were plated in opaque 384-well plates and transfected 24 hrs prior to the addition of formaldehyde or PBS (control/untreated). Cells were then grown for an additional 5 days prior to determining relative cell number using an Envision Multilabel detector/plate reader (Perkin Elmer). Luminescence values representing CTG reagent alone (blank) were subtracted from all wells to establish final luminescence values. A non-targeting universal siRNA negative control (MISSION siRNA SIC001 Universal Control #1) was used to monitor off-target effects, with results standardized as percent proliferation of siRNA-transfected compared to mock-transfected wells on the same plate. Compound additions were performed using peri-pumps as opposed to capillary pins. Pumps had the advantage of reproducibility and rapid delivery of 5 μL sample volumes, which minimized cell exposure times and led to reproducible signal intensities.

### RNAi screen data analyses

The means (*μ_t_*) for treated cell proliferation suppression were calculated from three replicate cultures for each gene and dose across each cell line. The mean for each dose treatment (*μ_t_*) was then normalized to the untreated mean (*μ_u_*) to calculate normalized proliferation suppression or mean (*μ_n_*).

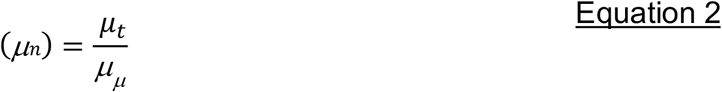

Normalized data were then used to calculate area under the curve (AUC) using GraphPad Prism 7 software (La Jolla, CA, USA). Z scores for AUC were calculated using the equation below, where *μ_a_* is the mean for the AUCs and *σ_a_* standard deviation for the AUCs.

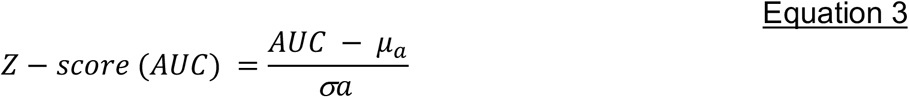

In each cell line, all genes with Z-scores ≤ 0 were considered sensitive. This cut-off score was chosen because we used an siRNA library already highly enriched for genes known to modulate cellular response to genotoxic agents, and for which sensitivities were expected to be less variable across genes. Highly sensitive genes were classified as genes with Z-score ≤ −1.0 for each cell line, while protective genes had Z score ≥ +2.0.

Hierarchical clustering with complete linkage and Euclidean distance was used to identify concordant proliferation suppression results among genes across cell lines. This analysis was performed on a 98-gene matrix where we had high quality relative viability measurements across all cell lines with no missing data. Based on visual examination of the resulting heatmap, we selected six gene clusters and used the ‘cutree’ R function to generate groups of genes informed by dendrogram height. For each cluster, we applied the multiple protein search query in String DB (Version 10.5) to assess the extent of protein-protein interactions (PPI) within these gene clusters (55), and used the STRING Analysis module to assess genome-wide KEGG pathway enrichment within clusters.

### Data Availability

Exome sequencing data are deposited in the Sequence Read Archive (SRA), accession number SRP131620.

## RESULTS

### Genes that suppress cell proliferation following formaldehyde exposure

We first determined formaldehyde dose-dependent cell proliferation suppression curves for the human cell lines GM00639, SW480, and U-2 OS. Suppression was assessed after five days of continuous formaldehyde exposure. All three cell lines displayed similar formaldehyde dose-dependent proliferation inhibition curves (Figure 1A), with similar formaldehyde doses leading to a 50% decrease in proliferation (expressed as GI_50_) (Figure 1B). Among the three lines, SW480 was the least and GM00639 the most sensitive to proliferation suppression at a specific formaldehyde concentration.

**Figure 1.**
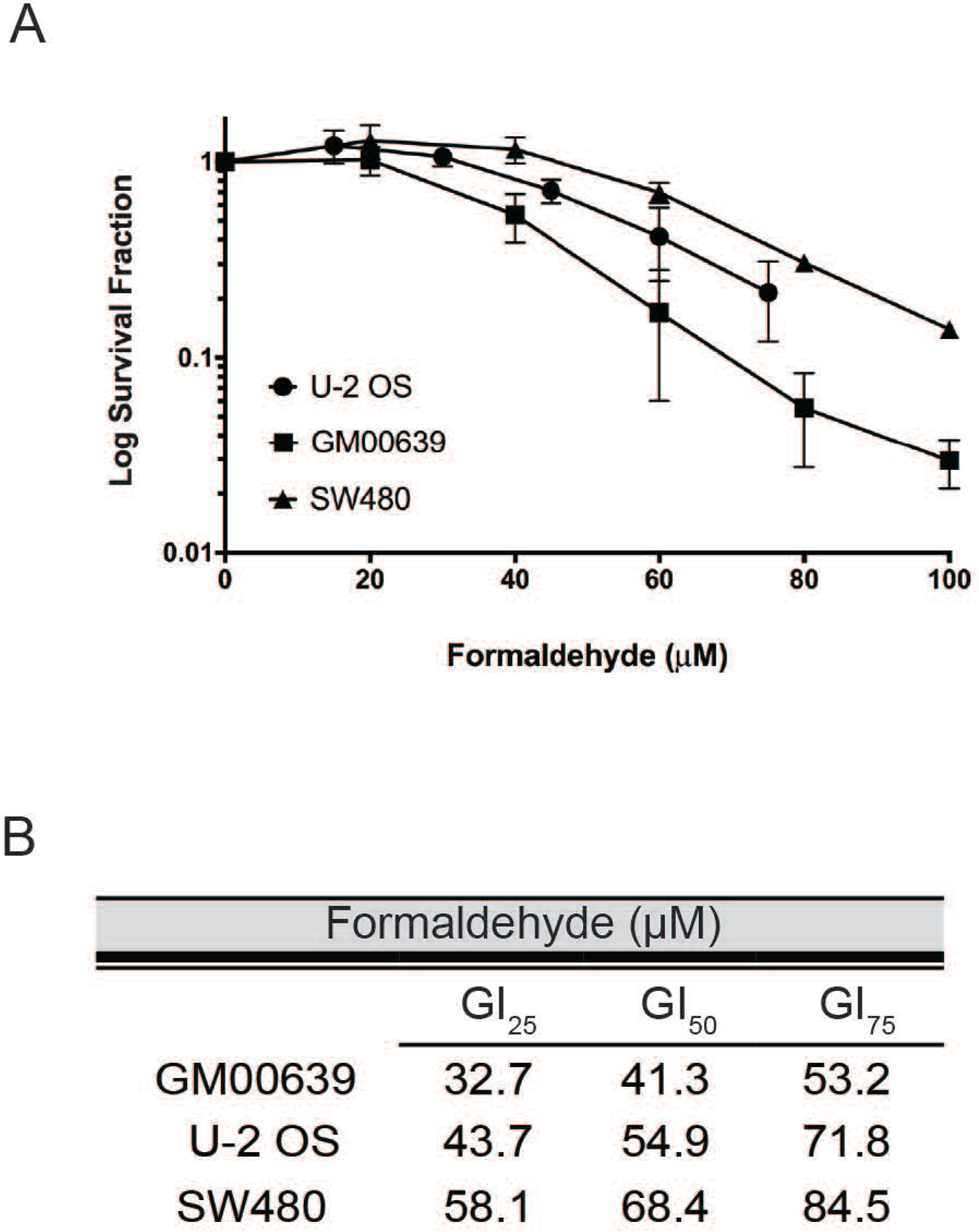
GM00639, U-2 OS, and SW480 have similar dose response curves following chronic formaldehyde exposure. (A) Dose response curves and (B) GI25-75 doses for each cell line following continuous formaldehyde exposure over 5 days.

We transfected each cell line, 24 h prior to formaldehyde exposure with our pooled siRNA library (54). Cell proliferation was assessed by CTG assay after 5 days of continuous formaldehyde exposure. A total of 23 genes conferred sensitivity when depleted across all three cell lines (Figure 2, Supplementary Table S1). These 23 genes represent pathways that may plausibly limit formaldehyde toxicity by promoting the repair of formaldehyde-induced DNA damage. Of note, no individual gene when depleted was either highly sensitizing (Z-score ≤ −1) or protective (Z-score ≥ +2) (Supplementary Figure S1A and B, respectively and Supplementary Table S1).

**Figure 2.**
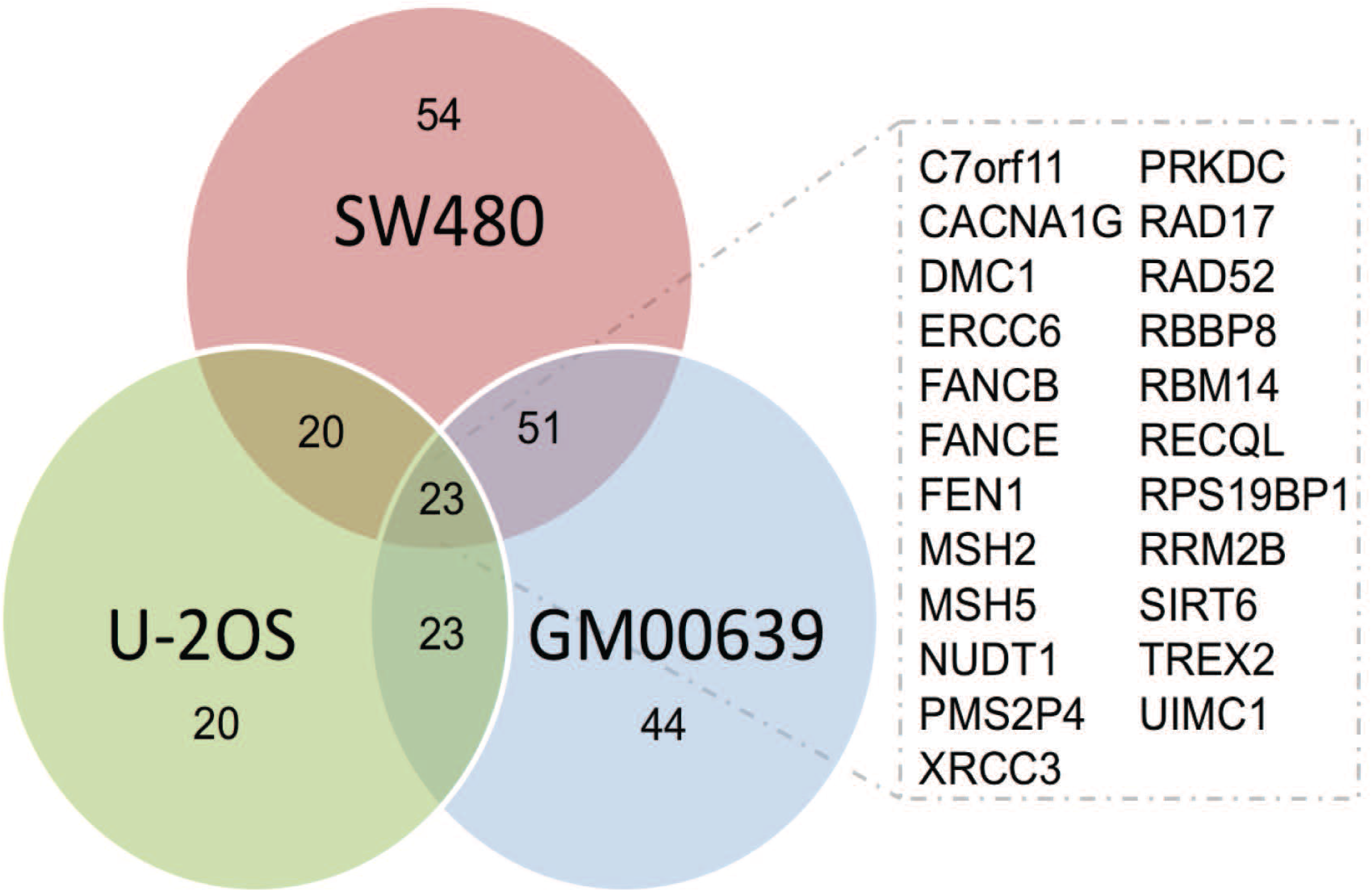
Twenty-three genes sensitize three human cell lines to formaldehyde. Venn diagram of the number of sensitizing genes (Z-score ≤ 0) in all three cell lines. Inset: List of 23 genes that were sensitizing in all three cell lines.

### Genomic alterations in the cell lines do not contribute to formaldehyde sensitivity

We analyzed non-synonymous mutations across cell lines to determine whether cell line-specific variability in the response to formaldehyde might be explained by cell line-specific genetic variation (Figure 3A and Supplementary Table S2). All three cell lines shared mutations in 2 genes (*PKHD1L1, TTN*) that were not included in our siRNA library. According to the ExAC database, both PKHD1L1 and TTN fall in the top 5% (z-score < −1.7) of genes over-represented for synonymous variation. Both genes also have loss-of-function probabilities of 0. These results indicate that PKHD1L1 and TTN are tolerant to variation, and suggests it is unlikely that these shared variant genes alter cell fitness (51).

**Figure 3.**
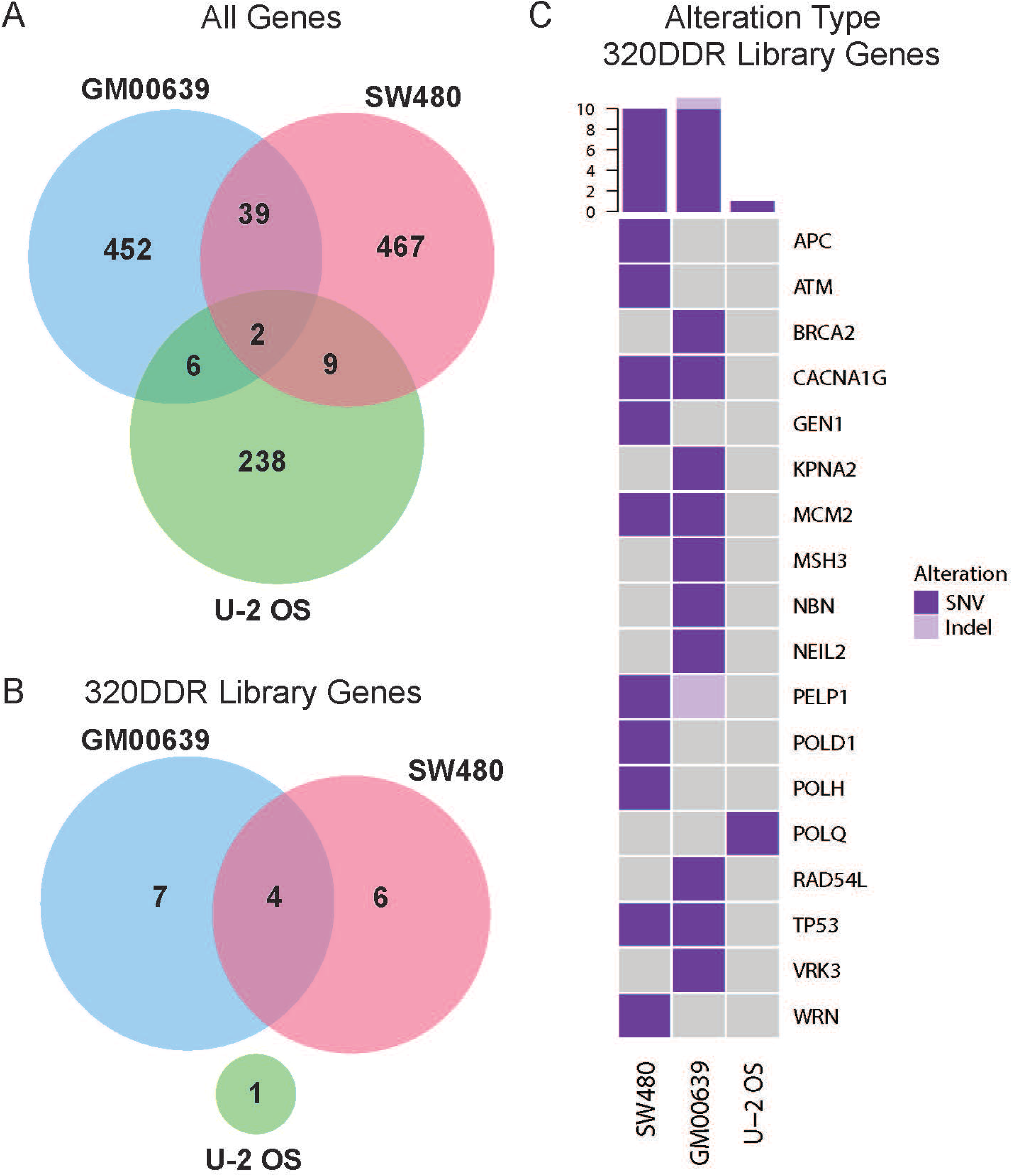
Prevalence of mutations in GM00639, SW480 and U-2 OS cell lines. Venn diagrams depicting the number of genes with nonsynonymous mutations in (A) All genes, versus (B) genes included in the ‘320 DDR’ Library across cell lines. Mutations include single nucleotide variants (SNVs) and small insertions and deletions (indels). (C) Oncoprint visualizing mutations in 320 DDR Library genes where each row represents a gene and each column a cell line (SW480, GM00639, or U-2 OS). Colors indicate a true event, *i.e*., a gene that is mutated in a given cell line. Gray indicates that no alteration was observed. The histogram (top) summarizes the number of genes affected for a given cell line.

No mutations were identified in siRNA target genes in all three lines (Figure 3B). However, 18 (or 6%) siRNA library-targeted genes were mutated in one cell line (Figure 3C) with no significant enrichment in specific DDR pathways (data not shown). These gene alterations may reflect a combination of donor-specific germline, or in the case of SW480 and U-2 OS cells somatic variants in the tumors from which these two lines were isolated. Thus, no common genomic alterations in siRNA-targeted genes explain the formaldehyde sensitivity or resistance.

Analysis of copy number variation on two of the three cell lines represented in the CCLE identified 269 genes in regions of copy number gain and 111 genes in regions of copy number loss shared between the SW480 and U-2 OS cell lines (Figure S2A-B, Table S3). Thirty-three (10%) genes targeted in the 320 DDR library were in regions of significant copy number variation (Figure S2C). We found a modest enrichment for amplified genes in cytokinesis pathway genes (*CETN2, KIF4A*) and for deleted genes in chromatin modifiers (*ATRX, CHAF1B*) in U-2 OS cells (Fisher’s exact test, p-value < 0.05). Apart from these associations, we identified no additional pathway enrichment for alterations in siRNA-targeted genes compared to all other genes assessed across all three cell lines. Of note, there was no enrichment in genomic alterations, either non-synonymous mutations or copy number changes, in genes known to modulate formaldehyde response when depleted versus control siRNAs. These results argue that genomic alterations in genes included in our 320 DDR library are not strong drivers of cell line-specific differences in formaldehyde response.

### Formaldehyde response pathways identified across cell lines

We extended our analyses by mapping genes targeted by our 320 DDR siRNA library to functional pathways. These were enriched for DNA metabolism and DNA damage response, reflecting the initial design of the 320 DDR library (54). Twenty-two functional gene groups were identified for siRNA library-targeted genes by combining Gene Ontology (GO) consortium terms, Reactome pathway data, and manual literature searches (Figure 4A and Table S4). Genes were assigned to multiple functional groups or pathways when appropriate (Figure 4B). Genes that when depleted sensitized cells to formaldehyde (*i.e*., Z-score ≤ 0) were significantly associated with eight functional pathways: HR, DSB Repair, Chromatin Modification, Cell Cycle, DNA Damage Checkpoints, Response to Oxidative Stress, Response to IR, and DNA replication (Figure 4C). An additional quantitative analysis of these associations, performed by bootstrapping with replacement, identified four functional pathways that were significantly over-represented among the sensitizing gene set: HR, DSB Repair, Response to IR, and DNA replication (Figure 4D).

**Figure 4.**
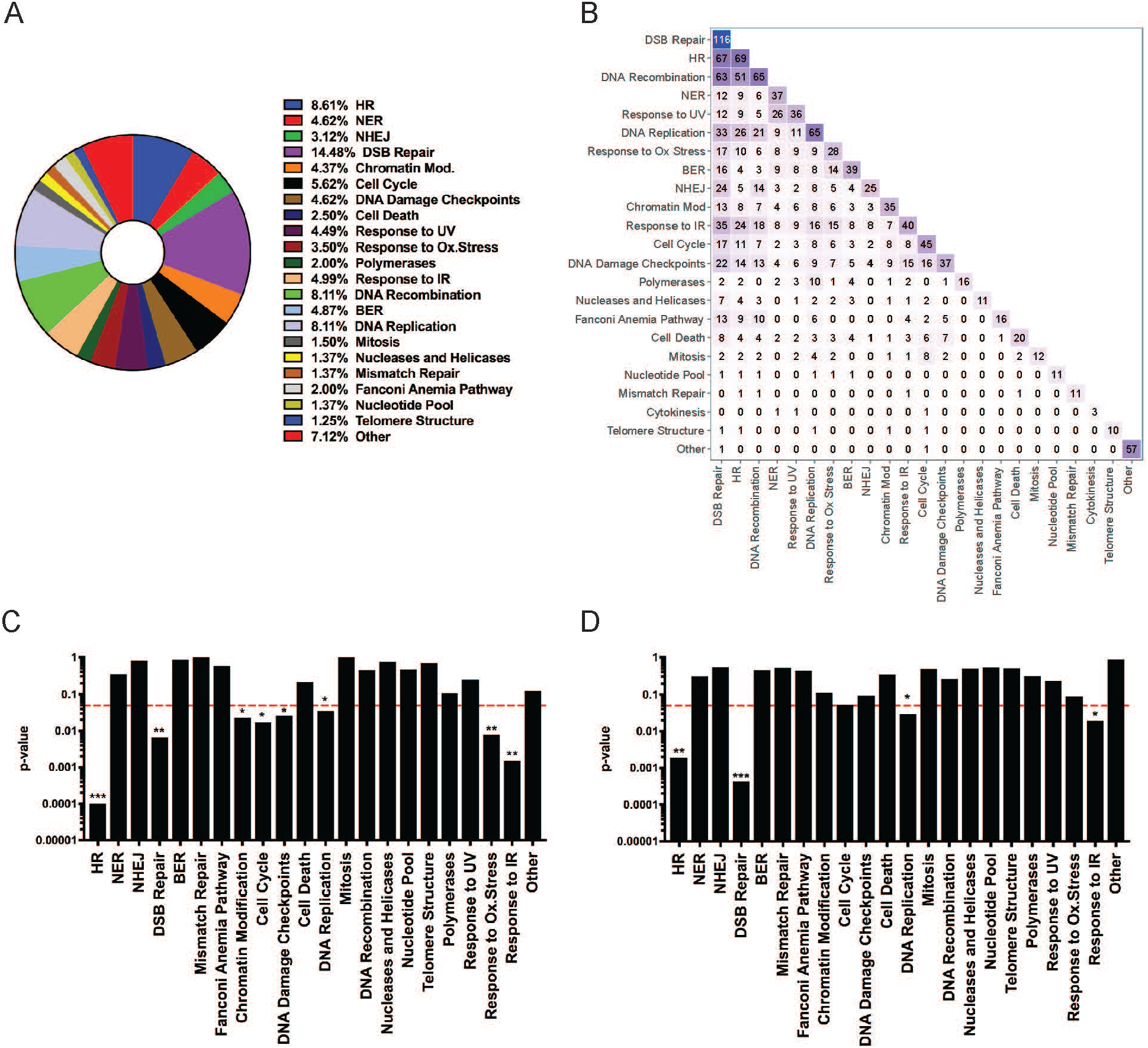
Enrichment analysis identifies pathways that mitigate proliferation suppression following formaldehyde exposure. (A) Fractional representation of genes included in 22 functional pathways that are represented by one or more genes targeted by our 320 DDR siRNA library. Genes present in more than one pathway were counted multiple times for these percentages. Percentages are not disambiguated for genes annotated to multiple pathways. (B) Heatmap summarizing the number of shared genes between annotated pathways. The diagonal represents the size of a given pathway annotation. Barplots depicting statistical significance (y-axis) of pathway enrichment testing by (C) Fisher’s exact test, and (D) bootstrapping with replacement with 1,000 iterations for the set of annotated pathways (x-axis). The dotted red line represents a p-value ≤ 0.05, threshold for significance.

The 23 sensitizing gene depletions did not belong to a single cellular pathway and were not genomically altered in a way that directly contributed to formaldehyde sensitivity. Thus, we interrogated gene product interaction networks to identify system-level cellular processes that might mitigate formaldehyde sensitivity. Hierarchical clustering of relative cell viabilities across cell lines for varying doses of formaldehyde identified two main clusters. The high dose formaldehyde samples (doses 3 and 4) primarily formed one cluster and low dose samples (doses 1 and 2) formed another cluster (Figure 5A). Genes in cluster 1 and 2 appear to mediate formaldehyde sensitivity across all cell lines at high doses, except for the U-2 OS cell line. Clusters 3–6 contained genes that when depleted led to cell line-specific variation in formaldehyde sensitivity. Pathway enrichment analysis showed an enrichment of genes in clusters 1 and 2 in 3 of the same KEGG pathways: NER, HR, and BER. Each cluster also displayed unique pathway enrichments, e.g., mismatch repair in cluster 2, or cell cycle and basal transcription factors in cluster 6 (Figure 5B). This approach identifies a concordance across cell lines in the pathways, but not gene products, necessary to mitigate formaldehyde-induced proliferation suppression.

**Figure 5.**
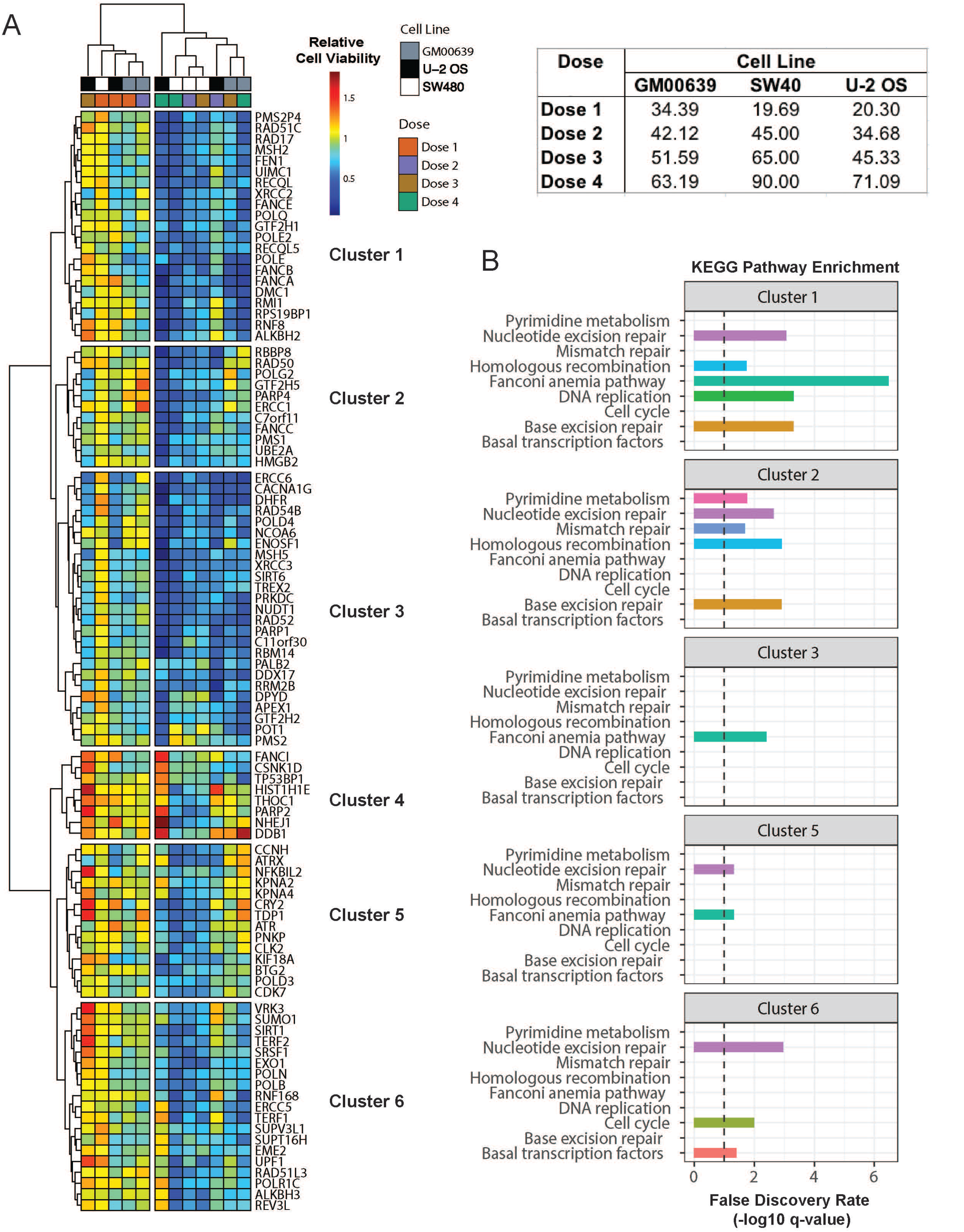
Relative cell proliferation reveals global and cell-line specific patterns of formaldehyde sensitivity. (A) Heatmap of relative cell proliferation suppression for a given formaldehyde dose across cell lines. Each map cell represents proliferation relative to an untreated control for a given gene and cell line. Hierarchical analyses of genes (rows) identified six clusters. Cell-line specific siRNA doses are shown in the table (top-right). (B) KEGG Pathway enrichment for gene clusters presented in A (FDR corrected p-values). Dotted lines represent an FDR threshold of 10%.

DNA damage repair pathways are highly conserved at both the functional and protein levels (56,57). Thus, we asked how well our human cell line results corresponded to our comparable essential gene yeast screen to identify genes that modulate cellular responses to formaldehyde (20). Among the 23 sensitizing genes we identified in all three human cell lines, 17 have a yeast homolog, including one which is essential and thus was not represented in our yeast deletion strain library. Of the 16 remaining genes from the human DDR screen, 9 also conferred formaldehyde sensitivity in our yeast deletion strain screen and mapped to comparable functional pathways: HR, DSB repair, DNA replication, DNA damage checkpoints, and cell cycle regulation (Table 1). Although deletion of *RAD57* did not sensitize cells to formaldehyde in our yeast screen, a comparable screen done in diploid yeast demonstrated that *RAD57* was required for formaldehyde tolerance (22). Together, these results identify several important functional pathways that strongly modulate formaldehyde toxicity across different functional pathways and eukaryotic species.

**Table 1.** Conserved genes and pathways that mitigate proliferation suppression following formaldehyde exposure. Human and *S. cerevisiae* gene names, along with the yeast systemic names are provided. A brief description of the gene pathway involvement, as well as the level of sensitivity in our previously published yeast screen, are noted (16).

| Human Gene | *S. cerevisiae* Gene | Systematic Name | Sensitivity | Description |
| --- | --- | --- | --- | --- |
| *CACNA1G* | *CCH1* | YGR217W | Not Sensitive | Calcium Voltage Channel |
| *DMC1* | *ECM30* | YLR436C | Sensitive | HR/DSB Repair/Mitosis |
| *ERCC6* | *RAD26* | YJR035W | Not Sensitive | NER/BER |
| *FANCB* | - | - | - | Fanconi Anemia Pathway |
| *FANCE* | - | - | - | Fanconi Anemia Pathway |
| *FEN1* | *RAD27* | YKL113C | Moderately | BER/DNA Replication |
| *MPLKIP* | *CDC5P* | YMR001C | Essential | Cell Cycle/ Mitosis |
| *MSH2* | *MSH2* | YOL090W | Moderately | Mismatch Repair |
| *MSH5* | *MSH5* | YDL154W | Not Sensitive | Mismatch Repair |
| *NUDT1* | *NPY1* | YGL067W | Not Sensitive | BER |
| *PMS2P4* | *PMS1* | YNL082W | Not Sensitive | Mismatch Repair |
| *PRKDC* | - | - | - | NHEJ/DSB Repair |
| *RAD17* | *RAD24* | YER173W | Sensitive | Cell Cycle/DNA Damage Checkpoint |
| *RAD52* | *RAD52* | YML032C | Sensitive | HR/DSB Repair/DNA Replication |
| *RBBP8* | *SAE2* | YGL175C | Not Sensitive | Cell Cycle/DNA Damage Checkpoint |
| *RBM14* | *PSP2* | YML017W | Sensitive | Mitochondrial mRNA Splicing |
| *RECQL* | *SGS1* | YMR190C | Sensitive | HR/DSB Repair/Replication |
| *RPS19BP1* | - | - | - | Ribosomal Subunit |
| *RRM2B* | *RNR4* | YGR180C | Sensitive | DNA Damage Checkpoint/ Replication |
| *SIRT6* | *SIR2* | YDL042C | Moderately | Chromatin Modification/DNA Replication |
| *TREX2* | - | - | - | HR/DSB Repair |
| *UIMC1* | - | - | - | HR/DSB Repair/Chromatin Modification |
| *XRCC3* | *RAD57* | YDR004W | Not Sensitive | HR/DSB Repair |

## DISCUSSION

Our analyses of genetic determinants of formaldehyde toxicity in human cells identified 23 genes and four functional pathways that modulate the response to chronic formaldehyde exposure across three human cell lines. We also identified 94 additional genes that conferred sensitivity when depleted in two cell lines, and an additional 118 genes that conferred sensitivity when depleted in a single cell line (Figure 2). The high fraction of siRNAs that modified the response of one or more cell lines to formaldehyde exposure likely reflects the design of the 320 DDR siRNA library which was focused on genes associated with DDR, replication, and repair (54). We identified four functional pathways that when perturbed strongly sensitized cells to formaldehyde: HR, DSB repair, DNA replication, and IR response. These pathways detect and respond to different types of macromolecular damage mediated by formaldehyde and are functionally interrelated in part due to the sharing of key proteins (Figure 4, Table S4).

One surprise in our screen data was that the Fanconi anemia (FA) pathway was not identified as a consistent contributor to mitigation of formaldehyde toxicity. However, siRNA-mediated depletion of 11 of the 22 FA complementation group genes included in the 320 DDR library did sensitize one or more cell lines to formaldehyde exposure (Figure 2, Table S1). This apparent lack of consistent Fanconi pathway-specific modulation across all three cell lines may reflect the redundant annotation of FANC proteins to additional pathways that mitigate formaldehyde toxicity and the presence of half of the currently recognized *FANC* genes as targets in our 320 DDR siRNA library (58) (Table S4). Genetic modifiers of FA pathway function may influence formaldehyde dose-dependent proliferation suppression in individual cell lines. Two groups of genetic modifiers with the potential to modify *FANC* gene function include the alcohol and aldehyde dehydrogenase (*ALDH* and *ADH*) gene families that catabolize both endogenous and exogenous formaldehyde and other aldehydes. Individual members of these gene families can strongly sensitize cells to aldehyde exposure and can promote disease progression in FA patients (29,59,60).

As part of our screen, we also identified several genes that conferred resistance to formaldehyde exposure in a specific cell line or genetic combination. Depletion of *ATM* was strongly protective in SW480 (Supplementary Table S1). This is reminiscent of other studies in which attenuation of the DNA damage response wherein ATM plays a key role blunted the toxicity of DNA damaging agents including IR (61,62). We also identified a protective effect of *DDB1* depletion in GM00639 and U-2 OS cells (Supplementary Table S1 and Supplementary Figure S1). DDB1 is the large subunit of the UV-damage DNA-binding protein complex (the UV-DDB complex) that participates in NER. It also participates in DCX (DDB1-CUL4-X-box) E3 ubiquitin-protein ligase complexes that may promote DNA repair by modifying individual proteins and chromatin (63). Of note, the depletion of *DDB2*, the protein heterodimer partner of DDB1, did not confer a similar protective effect (Supplementary Table S1), indicating that this protective effect may not depend directly on the UV-DDB complex.

Our results confirm and extend prior analyses of the genetic determinants of formaldehyde toxicity, and thus may have practical utility in at least two ways. First, the pathways we identified that sensitize cells to formaldehyde toxicity are all known to harbor substantial human genetic variation, including known pathogenic variants (64,65). As a result, our findings will help better define formaldehyde exposure-sensitized or resistant human genetic backgrounds and will further guide additional mechanistic analyses. The observation that some cancer cells can generate formaldehyde in response to chemotherapeutic exposure (11) raises a second intriguing idea: if this phenomenon is general, it may be possible to use a combination of cancer-targeting therapeutic agents and modifiers of the DNA damage response/repair pathways to potentiate cancer cell killing.

These potentially exciting extensions also highlight additional variables that need to be addressed before we can confidently extrapolate from cell-based screening data to tissue or organ-level effects. These toxicity determinants include: target tissue or organ cell types and their cycling or mitotic activity, tissue-level detoxification/quenching pathways, and the differential expression of genes that modify cellular responses following formaldehyde exposure. For example, the DNA damage response can vary greatly across tissue types and species as a function of tissue-specific gene expression and germline or somatic genetic variation (66–68). Our results also highlight the importance of performing studies in cell types relevant to the mode of exposure and cautions against making summary gene-specific statements in toxicity studies as gene expression, cell and tissue types, and both germline and acquired mutations can modulate cellular response. At the organismal level, both formaldehyde dose and route(s) of exposure will further strongly determine the responses to, and outcomes of, formaldehyde exposure. Once recognized and better understood, these potential sources of variability will provide a sound basis for performing genotoxic studies or for improving exposure and occupational hazard guidelines.

In summary, our results demonstrate the ability of systematic genetic screens to identify functionally important genes and pathways that modulate the response of human cells to formaldehyde exposure. Additionally, the genes and pathways identified in this study demonstrate a partial conservation of the functional pathways that mitigate formaldehyde toxicity across eukaryotes. Furthermore, our data reveal additional ways in which to predict, mitigate, and understand the biological consequences of human formaldehyde exposure.

## DATA AVAILABILITY

Raw exome sequencing data generated from the cell line GM00639 (CC1 clonal derivative) have been deposited in the Sequencing Read Archive SRA Accession: SRP131620.

## SUPPLEMENTARY DATA

Supplementary Data are available at NAR online.

## ACKNOWLEDGEMENTS

The authors are grateful to members of the McCullough lab, R. Stephen Lloyd, and Lloyd lab members for helpful discussions, James Annis of the Quellos HTS Facility for assistance in conducting the siRNA screen, and Dr. Laura Heiser for valuable insights regarding the data analyses.

## SUPPLEMENTARY TABLES AND FIGURES

**Supplementary Figure S1.**
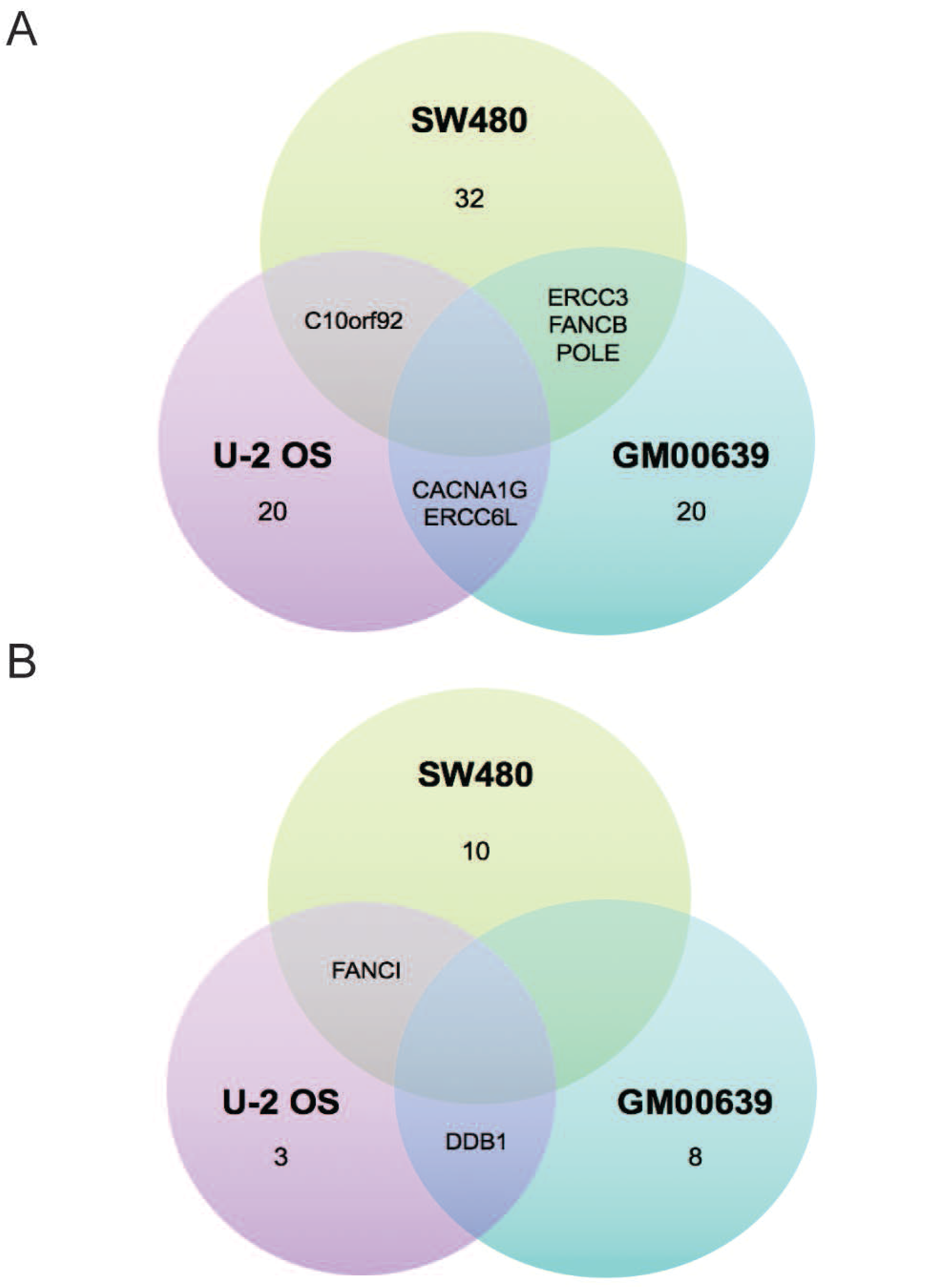
No gene was highly sensitizing or protective across all human cell lines. Venn diagrams summarize (A) the number of genes that were highly sensitizing (Z-score ≤ −1.0) to formaldehyde exposure for each cell line, and (B) the number or designation of genes that were protective following formaldehyde exposure, resulting in greater cell proliferation (Z-score ≥ +2.0) in each cell line.

**Supplementary Figure S2.**
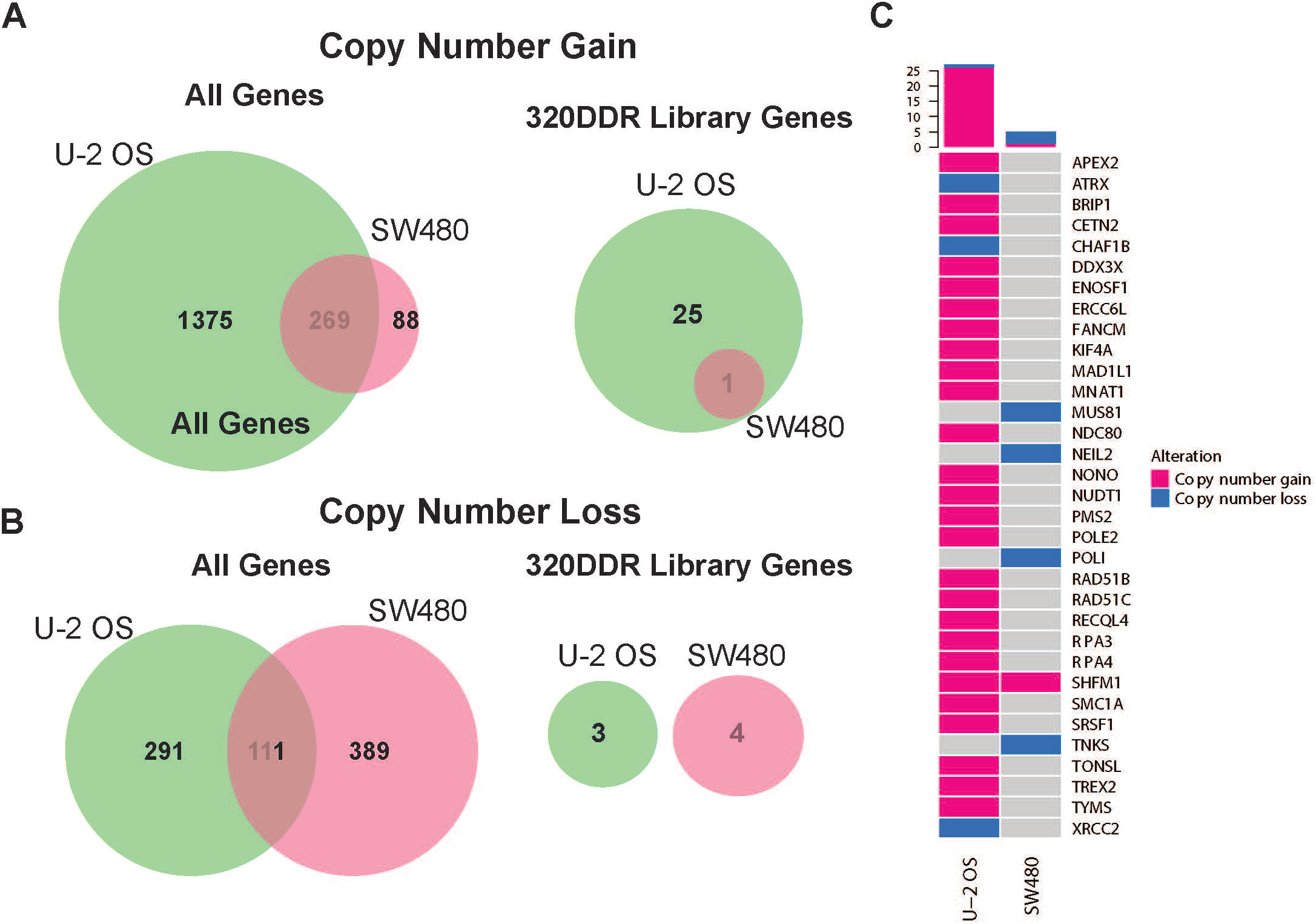
Copy number alterations do not influence formaldehyde sensitivity. Venn diagrams depicting the number of genes in regions of (A) copy number gain or (B) copy number loss in SW480 and U-2 OS cell lines for all genes (left) or 320 DDR Library genes (right). (C) Oncoprint visualizing copy number events in the 320 DDR Library genes for SW480 and U-2 OS cell lines. Each row represents a gene and each column a cell line (U-2 OS or SW480). Colors indicate a true event, *i.e*., a gene is in a region of copy number gain or loss for a given cell line. Gray indicates that no alteration was observed. The histogram (top) summarizes the number of genes affected for a given cell line.

**Supplementary Table S1. Cell line-specific Z-score tabs with indication of sensitization status in other two cell lines, rank-ordered by Z-score**. Sensitization status is determined using the Z-score threshold for highly sensitive (Z-score ≤ −1).

**Supplementary Table S2. Mutations in 320 DDR library genes for GM00639, SW480, and U-2 OS cell lines**. Annotated base-level variant calls in 320 DDR library genes.

**Supplementary Table S3. Copy number estimates in 320 DDR library genes for SW480 and U-2 OS cell lines**. Tabs indicate gene level copy number estimates for genes included in the 320 DDR siRNA library where copy number is higher (amplifications) or lower (deletions) than diploid, as indicated by mean segment log2 values.

**Supplementary Table S4. Gene Ontology and manual curations of gene pathway designation**. Genes were assigned to DDR pathway(s) using GO terms and manual curation of the literature. Shaded boxes indicate a positive pathway assignment.

